# Population history and the differential consequences of inbreeding and outcrossing in a plant metapoplation

**DOI:** 10.1101/2020.11.24.386946

**Authors:** Peter D. Fields, Gretchen Arnold, Joel M. Kniskern, Douglas R. Taylor

## Abstract

The phenotypic consequences of inbreeding typically result in a fitness decline proportional to the increase in the inbreeding coefficient, *F*. This basic assumption of a predictable, inverse relationship between fitness and *F* has been questioned by a number of empirical studies. We explored the relationship between population history and inbreeding in a metapopulation of the plant *Silene latifolia*, for which long-term data are available for the historical size and spatial distribution of hundreds of local demes. We used a population genetic analysis to estimate gene flow and bi-parental inbreeding (*F*_IS_) in demes with different histories of spatial isolation. A controlled crossing experiment examined whether the effect of inbreeding and outcrossing on fitness-related traits varied with different histories of population size and isolation. Historically isolated demes experienced less gene flow and an increase in *F*_IS_, as well as significant inbreeding advantage and outbreeding depression for traits expressed early in life. The causes of variation in the *F-*fitness relationship among populations will include variance in the distribution of deleterious recessive alleles driven by aspects of population history, including population size, founder effects, gene flow, bi-parental inbreeding, and opportunities for the purging of genetic load. Our findings show that isolation and historical variation in population size likely contribute substantial variation in past inbreeding and the consequences of future inbreeding across the metapopulation.

## INTRODUCTION

A major field of study in evolutionary biology attempts to understand the basis and consequences of different types of reproduction, which are amazingly diverse, especially in plants (Charlesworth, 2006). Recent comparative studies have suggested that the type of mating system can have profound effects on a species, including the species’ ability to persist (Heilbuth, 2000) and to both increase and maintain its range (Grossenbacher, *et al*., 2015). Numerous theories exist in order to transition these macro-scale patterns to micro-evolutionary processes, in particular focusing on the role of mating systems in generating variation in the distribution and magnitude of genetic load (Charlesworth and Willis, 2009). Less considered is the feedback between population processes, spatial distribution, and the ramifications of particular mating systems (Whitlock, 2002).

Natural populations of many species are subdivided into a number of smaller populations and interconnected by migration. The theory of geographically structured populations (or metapopulations) has shown that spatial structure can have profound effects on the distribution and magnitude of genetic load (Whitlock, 2002; Agrawal, 2010; Agrawal and Whitlock, 2012). Consider the fate of a deleterious recessive allele in geographically structured populations of obligate out-crossing species. Recessive alleles will be more exposed to selection because they experience higher relative frequency and homozygosity in a subset of demes. This results in inbreeding depression, which can be defined as the decline in fitness (or some trait related to fitness) associated with an increase in the inbreeding coefficient, *F*, over the short term, and perhaps a reduced genetic load (via more efficient purging of deleterious recessives) over the longer term (Thrall, *et al*., 1998; Keller and Waller, 2002; Whitlock, 2002). The genetic basis of inbreeding depression is caused either by the increased homozygosity of deleterious recessive alleles, or by decreasing relative frequency of heterozygotes at overdominant loci. Most research to date suggests that inbreeding depression is mainly caused by the presence of recessive deleterious mutations (Charlesworth and Willis, 2009). If geographic population structure is severe, fixation of deleterious recessive alleles may occur, a process that can only be reversed by inter-demic processes such as genetic rescue via migration (Willi and Fischer, 2005; Willi, Van Buskirk and Fischer, 2005). More generally, geographic population structure influences the outcome of natural selection whenever individuals interact ecologically or behaviorally with a local subset of conspecifics (McCauley, 1994). More generally, these aforementioned dynamics may be present or even more severe for species for which self-fertilization may also be possible (Escobar, Nicot and David, 2008). A natural consequence of the aforementioned definition of inbreeding depression in the present of segregating genetic variation in recessive deleterious alleles is that as *F* decreases for a pair of individuals resulting progeny, which may result from either among family or among population crosses, show higher fitness on average, or heterosis (Charlesworth and Willis, 2009). Importantly, in the presence of population genetic structure and heterosis rates of effective gene flow can be even higher than observed gene flow resulting from disproportionately larger fitness of offspring resulting from crosses between resident and migrant genotypes (Whitlock, Ingvarsson and Hatfield, 2000).

The basic assumption of a predictable, inverse relationship between fitness and *F* has been questioned by a number of recent empirical studies. In the plant *Ranunculus reptans* (creeping spearwort), inbred offspring may be equally fit or more fit relative to other individuals (Willi, *et al*., 2005). This study revealed substantial among-population variance in how fitness declines with increasing *F* (Willi, *et al*., 2005). A 5-generation serial inbreeding experiment in the angiosperm *Mimulus guttatus* showed that the relationship between total flower production (an assay for individual relative fitness) and degree of inbreeding varied significantly among populations and among individual lines, or families (Dudash, Carr and Fenster, 1997). In this experiment, the extinction of families that suffered higher levels of inbreeding, and the retention of families that suffered less from inbreeding, meant that the purging of genetic load could be accomplished more readily by selection among families, rather than selection among individuals within families. In *Physa acuta*, a freshwater snail, there was significant among-population, among-family, and among-population habitat type or class (river vs. pond) variance in the *F*-fitness relationship (Escobar, *et al*., 2008).

The causes of variation in the *F-*fitness relationship among populations, or among families within populations, must include some variance in the distribution of recessive, or nearly recessive, deleterious mutations. In a metapopulation, this will be caused by variance in population age, demographic history, historical population size, founder effect, gene flow, bi-parental inbreeding, and other past opportunities for the purging (or fixation) of deleterious recessive alleles (Zhou, Zhou and Pannell, 2010; Pannell and Fields, 2014). Perhaps the most considered parameter that will effect both the fixation of genetic load as well as its purging is effective population size, or *N*_E_, which in turn will be determined by metapopulation dynamics more generally. More specifically, very small population size can result in the fixation of deleterious alleles (whether recessive or otherwise) due to selection being less effective (Charlesworth, 2009). Concomitantly, as *N*_E_ increases selection should become more effective at removing individuals carrying larger genetic load, so called purging, though empirical evidence of this later dynamic is less clear (Byers and Waller, 1999; Charlesworth, 2009). In a metapopulation, these and related processes can occur at very local scales, and may combine to affect the level of inbreeding, inbreeding depression and opportunities for purging in an otherwise random-mating population (Whitlock, 2002).

Metapopulation dynamics may also generate local variation in phenotypic evolution that may in turn feed back to affect metapopulation processes such as gene flow, and population size and growth. Using the long-term study of a Finnish Glanville butterfly (*Melitaea cinxia*) metapopulation, Wheat et al. (2011) showed that females collected from newly colonized sites exhibited higher expression of abdomen genes involved in egg provisioning and thorax genes involved in the maintenance of flight muscle proteins, traits that might affect future gene flow or population size. If populations are segregating combinations of epistatically interacting loci and epistasis underlies reproductive isolation, then it follows that local colonization events may contribute to reproductive isolation. Matute (2013) conducted an experimental evolution study with *Drosophila yakuba*; one thousand replicate lines were forced through a genetic bottleneck, not unlike what might occur in metapopulations with extinction and recolonization in nature. While the most common outcome of the bottleneck and subsequent inbreeding was extinction, a number of lines persisted, simultaneously exhibiting premating isolation when crosses were made between other similarly bottlenecked lines. It is not clear how important these effects might be among newly colonized sites in nature, or the extent that such among population variance may be dissolved through the action of subsequent migration (Guillaume and Whitlock, 2007). Thus, in a metapopulation, local demes can be thought of as having potentially distinct histories of inbreeding, genetic drift, gene flow, mutation, and adaptive evolution (Eppley and Pannell, 2009; Zhou, *et al*., 2010). Just how distinct these histories are, and how long they persist, is an empirical question. Corbett-Detig et al. (2013), using a large-panel of *D. melanogaster* recombinant inbred lines (RILs), describe the genomic footprint of this very sort of epistatic interaction underlying within-species reproductive isolation, and suggest that the requisite genetic variation will likely segregate in natural populations rather than requiring new mutation implicit to the Dobzhansky–Muller model (Dobzhansky, 1937, 1941, 1951; Muller, 1942). Importantly, the role of these epistatic interactions become clearer in the context of metapopulation dynamics, wherein selection can act on non-additive genetic variance that becomes available via the colonization process (Goodnight 1998). Additionally, by reflecting and documenting metapopulation processes, the aforementioned deviations from the inverse relationship between fitness and *F*, a better understanding of the actual genetic basis of both inbreeding and outbreeding effects become clearer, and thereby may not entirely be the consequence of dominance masking deleterious recessive alleles (Whitlock 2002).

Using the angiosperm *Silene latifolia*, Richards (2000) identified recently established small populations, analogous to the recently bottlenecked lines in Matute (2013), showing that these newly-established populations were suffering from inbreeding depression relative to larger more established populations, and found evidence of enhanced gene flow, or genetic rescue, from outside sources. Our focus was to project this process forward, to test whether the contrasting histories of local populations led to different evolutionary outcomes, specifically in the relationship between inbreeding and fitness. We identified chronically small, isolated populations and used a fine-scale, molecular population genetic analysis to determine the consequences of population isolation on the quantity of inter-population gene flow and bi-parental inbreeding. We also use a combination of greenhouse experiments and controlled crosses to test whether small, isolated populations show different responses to further inbreeding or outcrossing. The present study aims to show how local variation in population history (population size, degree of spatial isolation) will affect the phenotypic consequences of inbreeding and thereby reveal mechanisms causing variation in the *F*-fitness relationship.

## MATERIALS AND METHODS

### STUDY ORGANISM

*Silene latifolia* is a dioecious perennial plant that was introduced to the United States from Europe. It occurs primarily in the northeast, but also inhabits higher elevations further south. The populations in this study are distributed in patches along the roadsides and farmland of southwestern Virginia, in the vicinity of Mountain Lake Biological Station. This region has been the focus of a metapopulation census since 1988 (Antonovics, *et al*., 1994; Richards, 2000). Census records consist of approximate numbers of male and female plants along continuous stretches of roadside.

### GENOTYPE SAMPLING

In order to determine the proportion of bi-parental inbreeding and gene flow across a large proportion of the metapopulation with different levels of spatial isolation, we sampled plants from 77 spatially distinct populations (Antonovics, *et al*., 1994; Antonovics, Thrall and Jarosz, 1998; Antonovics, 2004), during peak flowering in the summer of 2008, spanning ∼1/3 of the focal metapopulation (Figure 1). We collected leaf tissue from every plant in the population, or up to 50 individuals in the largest populations, and stored the leaves on silica gel (Sigma). Genomic DNA was extracted and amplified following established microsatellite techniques for *S. latifolia*. DNA was extracted from leaf tissue using the method described in (Keller, *et al*., 2012). We genotyped each individual plant at between 16 and 19 microsatellite loci. Our microsatellites are derived from multiple sources (Juillet, *et al*., 2003; Teixeira and Bernasconi, 2007; Moccia, *et* al., 2009; Abdoullaye, *et al*., 2010). PCR amplification was conducted using published methods for each marker. Three to four PCR products of different loci were then pooled together and added to a loading buffer containing formamide and GENESCAN 400HD ROX size standard (Applied Biosystems). Following five minutes of denaturing at 95 °C, fluorescently labeled fragments were separated on an Applied Biosystems 3130 sequencer and analyzed with GENEMAPPER v3.0 software (Applied Biosystems). Alleles were binned using the software TANDEM (Matschiner and Salzburger, 2009).

**Figure 1.**
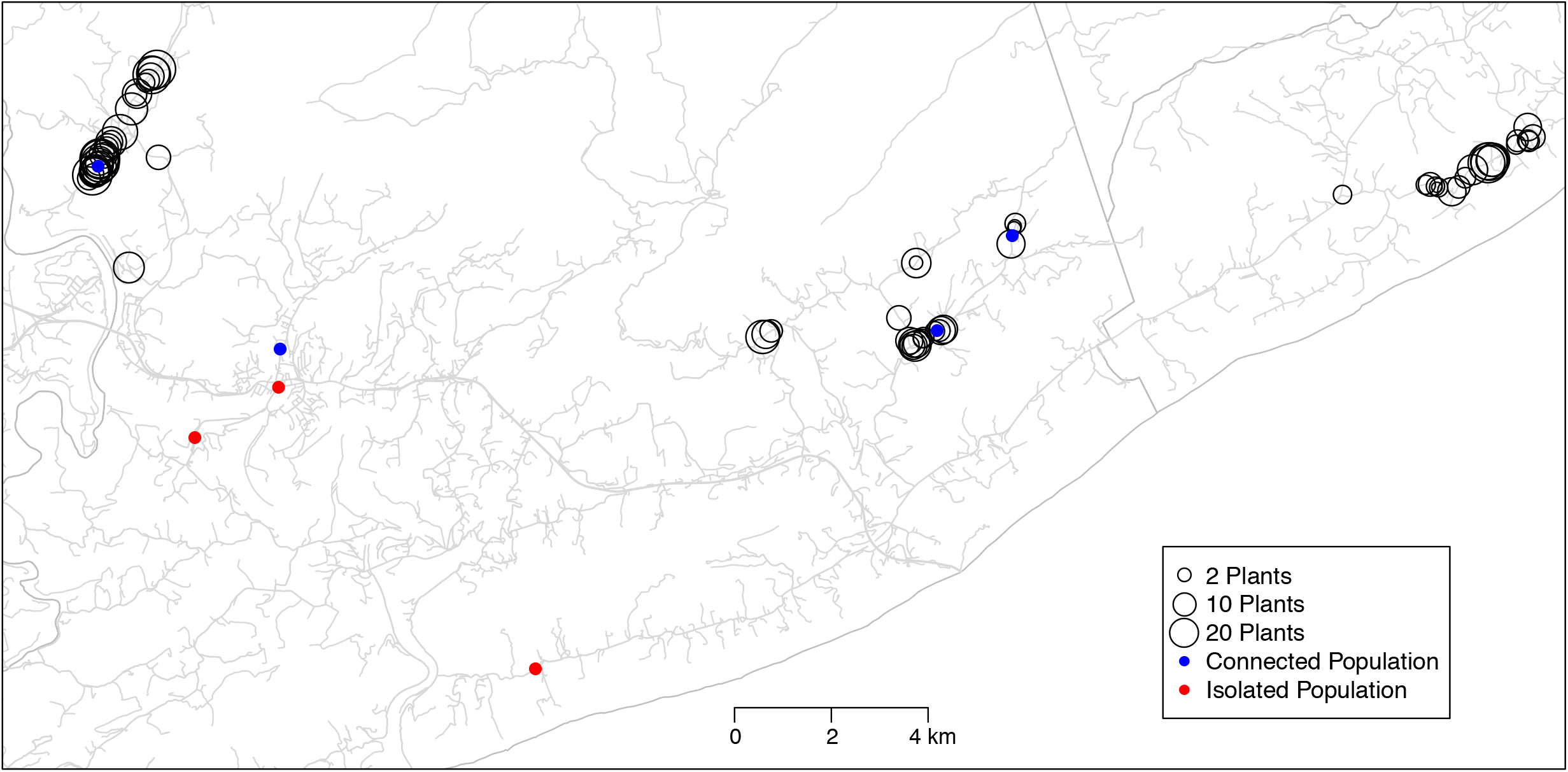
Map showing sampling locations for crossing design and population genetic analysis. Note, only populations used as part of the analysis are mapped. Other populations of *S. latifolia* exist in the sampled area.

### NATURAL POPULATIONS

To determine whether large, geographically central and small, geographically isolated populations differed in traits related to fitness, we recorded the number of seeds per capsule, seed mass, and germination percentage. We collected seed capsules from up to ten females in each of the small populations and three of the large populations (Figure 1). For each capsule, we counted the seeds and weighed them together to obtain the mean seed mass. From each capsule, we then planted groups of five seeds in each of five one-inch tubular pots that contained a standard, homogeneous soil mixture. Pots were arranged randomly in racks indoors and were watered daily. We recorded the number of days until the cotyledons emerged. To increase the reliability of our estimate of germination percentage, we also recorded the germination percentage of seeds from these six populations and two additional large populations. After the seeds had aged at least five weeks, we put 100 seeds from each capsule into petri plates lined with filter paper, and recorded the proportion that germinated.

### CROSSING EXPERIMENTS

For our controlled crosses, we confined this study to two types of populations: those that had been isolated for the entire duration of the census and those that have been consistently centrally located. Three small populations were used, ranging from approximately 10 to 30 individuals. Using 10-years of census data and correcting for sex ratio and among year variation in population size, we calculated the average demographic effective population size (*N*_*E*_) to be 8.7 for the small populations (Wright, 1938; Caballero, 1994; Ingvarsson, 1998). Population sizes from our metapopulation census data are generally underestimates relative to more detailed demographic studies we have done, but the relative size of populations is generally accurate. Two of the small populations were more than 1800m from the nearest neighboring patch of plants. The third small population was 360m from the nearest neighbor. Pollen flow in *S. latifolia* typically does not exceed 100m (Richards, Church and McCauley, 1999) and divergence in gene frequency occurs over a distance of approximately 150m or even less (McCauley, *et al*., 1996). Thus, the small populations in this study were expected to be relatively isolated. For large, centrally located populations, five populations that had well over 100 individuals for the past 10 years were selected (*N_E_*=105.8). One large population experienced a demographic bottleneck when numbers were reduced from 200 to 50 individuals. One large population is not within the metapopulation census, so nothing is known about its specific demographic history, except that it has been a large field with hundreds of plants since the beginning of the census (1988) (D.R. Taylor, pers. obs.).

To determine whether differences in early life-history traits between small, isolated and large, central populations were genetically based, and to test the predictions of the purging and drift models for small populations, we carried out a series of crosses in the greenhouse. If small, isolated populations had fixed deleterious recessive alleles through the overwhelming role of genetic drift, we expect plants from those populations to have lower fitness, with relatively little reduction in fitness with further inbreeding, but with higher fitness when plants are crossed among populations. If small, isolated populations have tended to purge deleterious recessive alleles, which given small population size is unlikely given theoretical expectations, we expect no overall reduction in plant fitness, little or no inbreeding depression, and little or no advantage to outcrossing (relative to larger populations). Rather, if the observed degree of genetic load is low in these populations, the colonization process itself must have selected for propagules with lower quantities of deleterious recessive alleles.

Three small, isolated populations and five large, central populations were used. We randomly selected a male and a female from up to ten families per population (some small populations had less than ten families). Each female was crossed with three males: 1) the male plant from her family (sib-mating), 2) a randomly selected non-sibling male from within her own population (random mating), and 3) a randomly selected male from another population used in the study (outcrossing). All pollinations were carried out with male and female flowers that had opened in the previous 24 hours. Seeds per capsule, mean seed mass, and germination was measured on the resulting fruits.

## DATA ANALYSIS

### POPULATION GENETIC ANALYSIS OF CONNECTIVITY AND BI-PARENTAL INBREEDING

There are a number of methods described in published literature that might be used to estimate population isolation. The majority of ecological studies have utilized a nearest neighbor/patch approach, or distance to multiple neighbors within a limited neighborhood of a focal patch (or buffer) (Moilanen and Nieminen, 2002). However, these simple measures have been shown to be poor predictors of important metapopulation dynamics such as colonization potential (Moilanen and Nieminen, 2002). Instead, we used a sum total of all pair-wise distances of a focal population to all other extant populations within an individual metapopulation section (each section is separated by >> 1 kilometer) (Gaggiotti, *et al*., 2009; Mora, *et al*., 2010). These pair-wise distances were calculated using a network constructed based upon the public roadway system, using ArcGIS (ESRI) Network Analyst tool. Given the mountain-valley geographic topology of the area, this network-based approach is more appropriate than standard Euclidean distances in order to predict likely routes of the predominant pollinators, noctuid moths. As such, larger isolation scores are indicative of a decrease in an individual population’s probability of receiving migrants, whether through seeds or pollen.

We calculated the population genetic summaries of genetic diversity, as well as estimates of genetic (sub)-structure via hierarchical *F*-statistics using the software GenoDive version 2.0b21 (Meirmans and Van Tienderen, 2004), with significant deviations from 0 assessed with 10,000 permutations and *α*= 0.05.

To assess how isolation affects migration among populations, we used the program BayesAss v. 3.03 (Wilson and Rannala, 2003). Like above, we analyzed each section separately, as the likelihood of migration from one section to the other is quite low. A total of three runs per section were done, each using 50,000,000 Markov Chain Monte Carlo (MCMC) iterations and a burn-in of 500,000 iterations, and a thinning interval of 100, each with a different starting seed. In order to obtain appropriate mixing conditions, as determined by acceptance rate, in the MCMC chain, we modified the allele frequency, inbreeding coefficient, and migration rate parameters as per the BayesAss v. 3.03 manual suggestion. Chain convergence was assessed using the program Tracer v. 1.5 (Rambaut and Drummond, 2009).

We tested for significant consequences of isolation on population genetic summaries and gene flow using Gaussian linear models. Linear modeling of the effects of isolation on these population genetic parameters were performed in R v. 2.15.3.(R Development Core Team, 2012).

### NATURAL POPULATIONS

Each trait was analyzed to determine whether mean values differed significantly for large and small populations in nature. Seed counts were normalized with a square-root transformation, and germination percentage was normalized with an arcsine transformation. These two traits were analyzed by ANOVA with the populations treated as random effects and nested within the size class of the population. The data on mean seed mass were distinctly bi-modal and were analyzed using Wilcoxon Rank-Sum tests.

### CROSSING EXPERIMENTS

To determine whether there was a genetic basis to the differences between population size classes in the field and assess the effects of further inbreeding/outcrossing, we used ANOVA to compare the trait means in the offspring of plants from large, central and small, isolated populations. The offspring were derived from our crossing scheme. We conducted a two-way ANOVA in order to determine the role of population type, crossing treatment, and their interaction. The ANOVA was a mixed model with population size class (large, central versus small, isolated) and population cross type as fixed effects and population (nested within size class) as a random effect. In order to test particular hypotheses about the origins of variation in the consequences of particular mating types (Escobar, *et al*., 2008), we conducted separate analyses for random within-population crosses, among population crosses, and sib-matings. The ANOVA was a mixed model with population size class (large versus small) as a fixed effect and population (nested within size class) as a random effect. To meet the assumptions of ANOVA, seed count was square root transformed and seed mass was log-transformed. The response variable for germination was the generalized logit of the ratio of the number of seeds that did not germinate to the number of seeds that did germinate.

When evaluating the fitness advantages/disadvantages associated with inbreeding and outcrossing, we made certain assumptions about how each trait was related to fitness. Fewer days to germination was assumed to positively affect fitness based upon assumptions of viability selection and seedling competition on perennial species (Baskin and Baskin, 1998). Relative fitness from inbreeding was calculated as 1-(value from sib cross / value from random within population cross). Relative fitness from outbreeding was calculated as 1-(value from outcross / value from random within population cross). All other traits were assumed to be positively correlated with fitness, where relative fitness from inbreeding=(value from sib cross / value from random within population cross)-1 and relative fitness from outbreeding=(value from outcross / value from random within population cross)-1. One-way ANOVA was used to determine the significance of the differences between the two relevant cross types on the trait.

## RESULTS

### GENETIC ANALYSIS OF CONNECTIVITY AND BI-PARENTAL INBREEDING

Individual microsatellites varied in the amount of population (sub)structure, though only two markers in two of the three sections showed non-significant *F*_ST_ values. The global *F*_IS_ values for the three separate sections of the metapopulation were 0.229, 0.272, 0.317, respectively, with a range of −0.008 and 0.64 for individual markers across metapopulation sections (Table 1; Suppl. Table 1). Populations varied in their level of genetic isolation. There was significant gene flow between populations with a global average of ∼28% individuals within populations being a recent migrant (Table 1; Suppl. Table 1).

**Table 1.** Global population genetic summaries. Variables are the number of alleles (*N*), the effective number of alleles (*E_N*), observed heterozygosity (*H*_*O*_), Expected heterozygosity (*H*_*E*_), and the inbreeding coefficient (*F*_IS_), global among-population allelic variation (*F*_IS_), and Jost’s measure of population differentiation (*D*_EST_).

| Locus | Section 2 |  |  |  |  |  |  | Section 6 |  |  |  |  |  |  | Section 9 |  |  |  |  |  |  |
| --- | --- | --- | --- | --- | --- | --- | --- | --- | --- | --- | --- | --- | --- | --- | --- | --- | --- | --- | --- | --- | --- |
|  | N | E_N | H<sub>O</sub> | H<sub>E</sub> | F<sub>IS</sub> | F<sub>ST</sub> | D<sub>EST</sub> | N | E_N | H<sub>O</sub> | H<sub>E</sub> | F<sub>IS</sub> | F<sub>ST</sub> | D<sub>EST</sub> | N | E_N | H<sub>O</sub> | H<sub>E</sub> | F<sub>IS</sub> | F<sub>ST</sub> | D<sub>EST</sub> |
| SL_eSSR01<sup>1</sup> | - | - | - | - | - | - | - | 7 | 1.818 | 0.224 | 0.488 | 0.542 | **0.114** | 0.129 | 6 | 1.722 | 0.161 | 0.452 | 0.644 | **0.099** | 0.094 |
| SL_eSSR04<sup>1</sup> | 10 | 2.282 | 0.577 | 0.584 | 0.012 | **0.05** | 0.076 | 7 | 2.399 | 0.524 | 0.62 | 0.154 | **0.05** | 0.089 | 6 | 2.475 | 0.519 | 0.629 | 0.174 | **0.105** | 0.206 |
| SL_eSSR06<sup>1</sup> | 18 | 3.762 | 0.699 | 0.765 | 0.086 | **0.041** | 0.145 | 14 | 3.333 | 0.503 | 0.751 | 0.33 | **0.069** | 0.233 | 13 | 3.108 | 0.39 | 0.725 | 0.462 | **0.1** | 0.304 |
| SL_eSSR09<sup>1</sup> | 12 | 1.407 | 0.287 | 0.301 | 0.049 | **0.037** | 0.017 | 7 | 1.65 | 0.393 | 0.416 | 0.055 | **0.073** | 0.059 | 9 | 1.584 | 0.382 | 0.386 | 0.01 | **0.091** | 0.065 |
| SL_eSSR12<sup>1</sup> | 20 | 3.296 | 0.724 | 0.724 | -0.001 | **0.044** | 0.123 | 14 | 3.557 | 0.674 | 0.762 | 0.116 | **0.117** | 0.444 | 15 | 4.093 | 0.702 | 0.795 | 0.117 | **0.087** | 0.381 |
| SL_eSSR16<sup>1</sup> | 8 | 1.569 | 0.334 | 0.378 | 0.117 | **0.099** | 0.068 | 4 | 2.245 | 0.583 | 0.584 | 0.002 | **0.073** | 0.116 | 8 | 2.248 | 0.537 | 0.583 | 0.079 | **0.141** | 0.238 |
| SL_eSSR17<sup>1</sup> | - | - | - | - | - | - | - | 13 | 2.747 | 0.608 | 0.674 | 0.098 | **0.106** | 0.256 | 14 | 2.66 | 0.556 | 0.658 | 0.154 | **0.069** | 0.147 |
| SL_eSSR20<sup>1</sup> | 11 | 1.161 | 0.121 | 0.145 | 0.168 | **0.065** | 0.012 | 3 | 1.057 | 0.056 | 0.057 | 0.016 | 0.028 | 0.002 | 3 | 1.048 | 0.038 | 0.049 | 0.218 | 0.039 | 0.002 |
| SL_eSSR22<sup>1</sup> | - | - | - | - | - | - | - | 10 | 2.817 | 0.561 | 0.686 | 0.183 | **0.149** | 0.403 | 11 | 3.167 | 0.501 | 0.726 | 0.31 | **0.084** | 0.253 |
| SL_eSSR27<sup>1</sup> | - | - | - | - | - | - | - | 9 | 2.462 | 0.58 | 0.628 | 0.077 | **0.167** | 0.355 | 11 | 2.568 | 0.595 | 0.641 | 0.071 | **0.128** | 0.271 |
| SL_eSSR28<sup>1</sup> | - | - | - | - | - | - | - | 6 | 1.79 | 0.469 | 0.465 | -0.008 | **0.22** | 0.257 | 6 | 1.353 | 0.24 | 0.275 | 0.126 | **0.245** | 0.127 |
| SL_eSSR29<sup>1</sup> | 25 | 5.488 | 0.835 | 0.85 | 0.018 | **0.054** | 0.332 | 19 | 3.871 | 0.715 | 0.785 | 0.09 | **0.103** | 0.441 | 18 | 3.683 | 0.7 | 0.765 | 0.085 | **0.083** | 0.305 |
| SL_eSSR30<sup>1</sup> | 21 | 3.407 | 0.65 | 0.737 | 0.119 | **0.056** | 0.17 | 21 | 2.683 | 0.345 | 0.679 | 0.492 | **0.111** | 0.278 | 18 | 2.705 | 0.281 | 0.678 | 0.585 | **0.083** | 0.197 |
| slat_18<sup>2</sup> | 10 | 2.837 | 0.418 | 0.683 | 0.388 | **0.058** | 0.137 | 9 | 2.045 | 0.234 | 0.556 | 0.578 | **0.208** | 0.346 | 11 | 2.522 | 0.276 | 0.649 | 0.575 | **0.119** | 0.259 |
| slat_32<sup>2</sup> | 9 | 2.709 | 0.29 | 0.67 | 0.568 | **0.137** | 0.331 | 7 | 2.271 | 0.364 | 0.602 | 0.396 | **0.159** | 0.301 | 9 | 2.384 | 0.34 | 0.62 | 0.451 | **0.181** | 0.375 |
| slat_33<sup>2</sup> | 3 | 1.128 | 0.061 | 0.12 | 0.49 | **0.237** | 0.044 | 2 | 1.515 | 0.184 | 0.368 | 0.502 | **0.265** | 0.221 | 3 | 1.211 | 0.105 | 0.186 | 0.434 | **0.597** | 0.35 |
| slat_48<sup>2</sup> | 6 | 1.253 | 0.087 | 0.214 | 0.593 | **0.099** | 0.031 | 5 | 1.766 | 0.194 | 0.472 | 0.589 | **0.108** | 0.113 | 5 | 1.666 | 0.233 | 0.427 | 0.455 | **0.139** | 0.125 |
| slat_72<sup>2</sup> | 12 | 2.858 | 0.349 | 0.688 | 0.493 | **0.048** | 0.116 | 12 | 2.545 | 0.326 | 0.657 | 0.504 | **0.097** | 0.217 | 16 | 2.86 | 0.282 | 0.7 | 0.596 | **0.171** | 0.499 |
| slat_85<sup>2</sup> | 15 | 2.085 | 0.311 | 0.55 | 0.434 | **0.101** | 0.142 | 16 | 1.829 | 0.284 | 0.489 | 0.418 | **0.156** | 0.185 | 16 | 1.618 | 0.232 | 0.408 | 0.432 | **0.194** | 0.172 |
| Sl_8<sup>3</sup> | 11 | 2.153 | 0.324 | 0.566 | 0.428 | **0.132** | 0.203 | - | - | - | - | - | - | - | - | - | - | - | - | - | - |
| A11<sup>4</sup> | 43 | 4.382 | 0.707 | 0.806 | 0.122 | **0.062** | 0.283 | - | - | - | - | - | - | - | - | - | - | - | - | - | - |
| Overall | 14.625 | 2.611 | 0.423 | 0.549 | 0.229 | **0.074** | 0.099 | 9.737 | 2.337 | 0.412 | 0.565 | 0.272 | **0.128** | 0.2 | 10.421 | 2.351 | 0.372 | 0.545 | 0.317 | **0.138** | 0.198 |

Linear models were used to test the effect of population spatial isolation on resident proportion (a BayesAssv3 based estimate of the proportion of a population that is non-migrant) and on multi-locus *F*_IS_ (in this case, an approximate estimate of bi-parental inbreeding), while controlling for population size as part of the model. The log of sum isolation had a highly significant effect on resident proportion (P-value < 0.001, adj. R^2^ = 0.4904) and multi-locus *F*_IS_ (P-value = 0.0114, adj. R^2^ 0.1083) (Figure 2).

**Figure 2.**
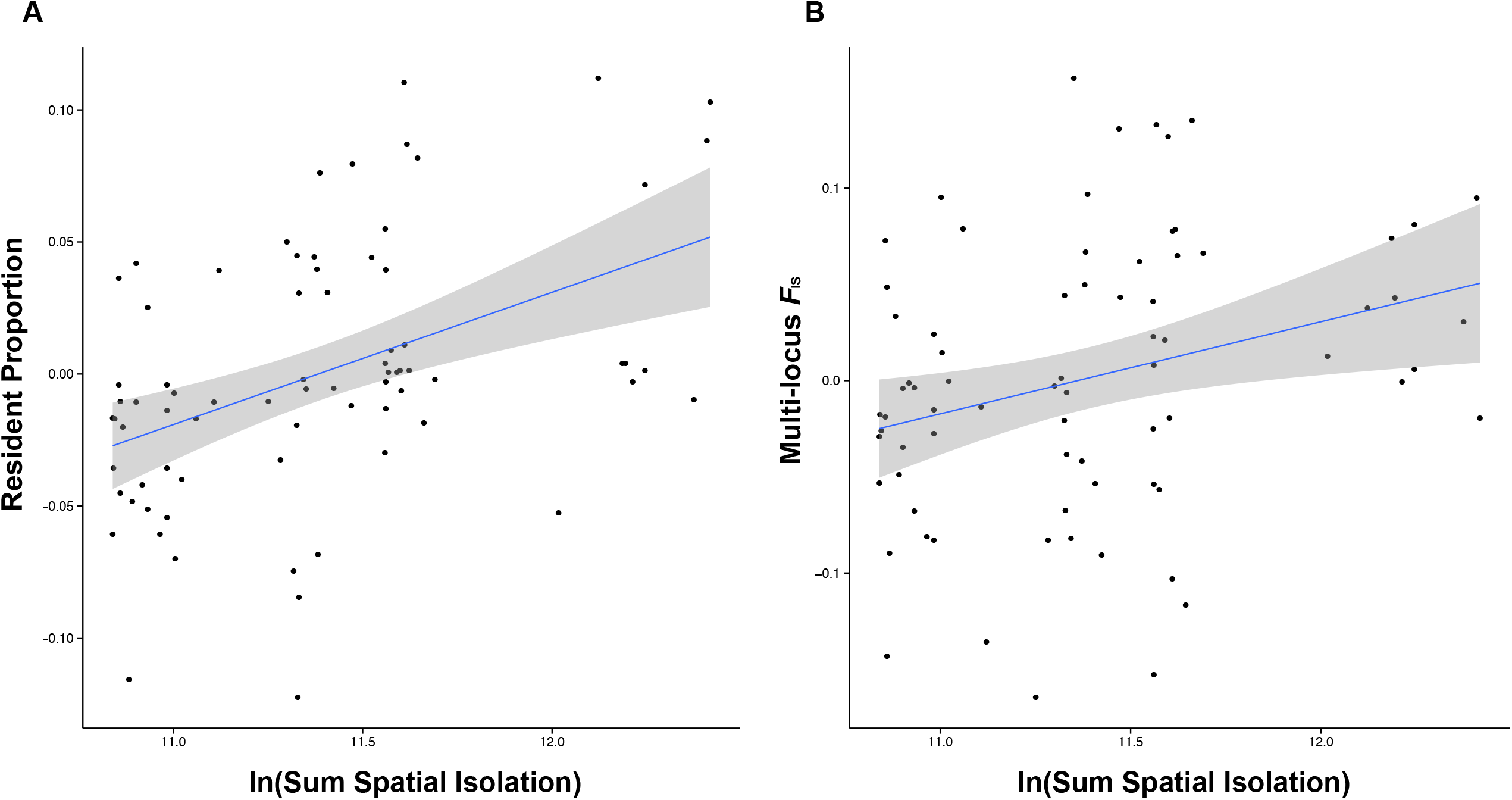
Consequences of spatial isolation on (A) resident proportion and (B) multi-locus *F*_IS_. The log of sum isolation had a significant effect on multi-locus *F*_IS_ (P-value = 0.0114, adj. R^2^ 0.1083) and resident proportion (P-value < 0.001, adj. R^2^ = 0.4904).

### NATURAL POPULATIONS

Measurements on plants in the field detected two significant differences between large and small populations in the means of three fitness-related traits (Table 2). Plants in large populations produced more seeds per capsule and larger seeds than plants in small populations.

**Table 2.** Means (and standard deviations) of seed and seedling traits of seeds collected from central and small, isolated populations in the field. Seed mass is the total weight of 50 seeds from a single capsule. P-values represent the significance of the difference in mean between large and small populations (see text for details).

| **Population Type** | **Seed Number** | **Seed Mass (g)** | **Germination Percentage** |
| --- | --- | --- | --- |
| Large, central | 277<br>(-28) | 0.054<br>(-0.005) | 69<br>(-15) |
| Small, isolated | 134<br>(-64) | 0.04<br>(-0.01) | 71<br>(-15) |
| **P-values** | **<0.0001** | **0.0041** | 0.2749 |

### GENETICS OF FIELD POPULATIONS

The results from our hand-pollinations showed that the difference in fitness were at least partially genetically determined. In particular, a significant interaction effect for seed mass (P < 0.005), and a nearly significant interaction for both seed number and germination percentage (P = 0.059 and P= 0.074, respectively) suggest a distinct role of population history in determining the consequences of particular mating types Table 3. Individual ANOVAs allowed us to compare the field measured values and the green house measured values, as well as test particular hypotheses about the effect of population history in generating different consequences of inbreeding and outbreeding. We found high similarity between field and green house measured traits, in particular seed mass, between large, central and small, isolated populations, and were maintained in within-population crosses in the greenhouse (Tables 2, 4). Seed number and germination showed no significant differences among population size classes.

**Table 3.**
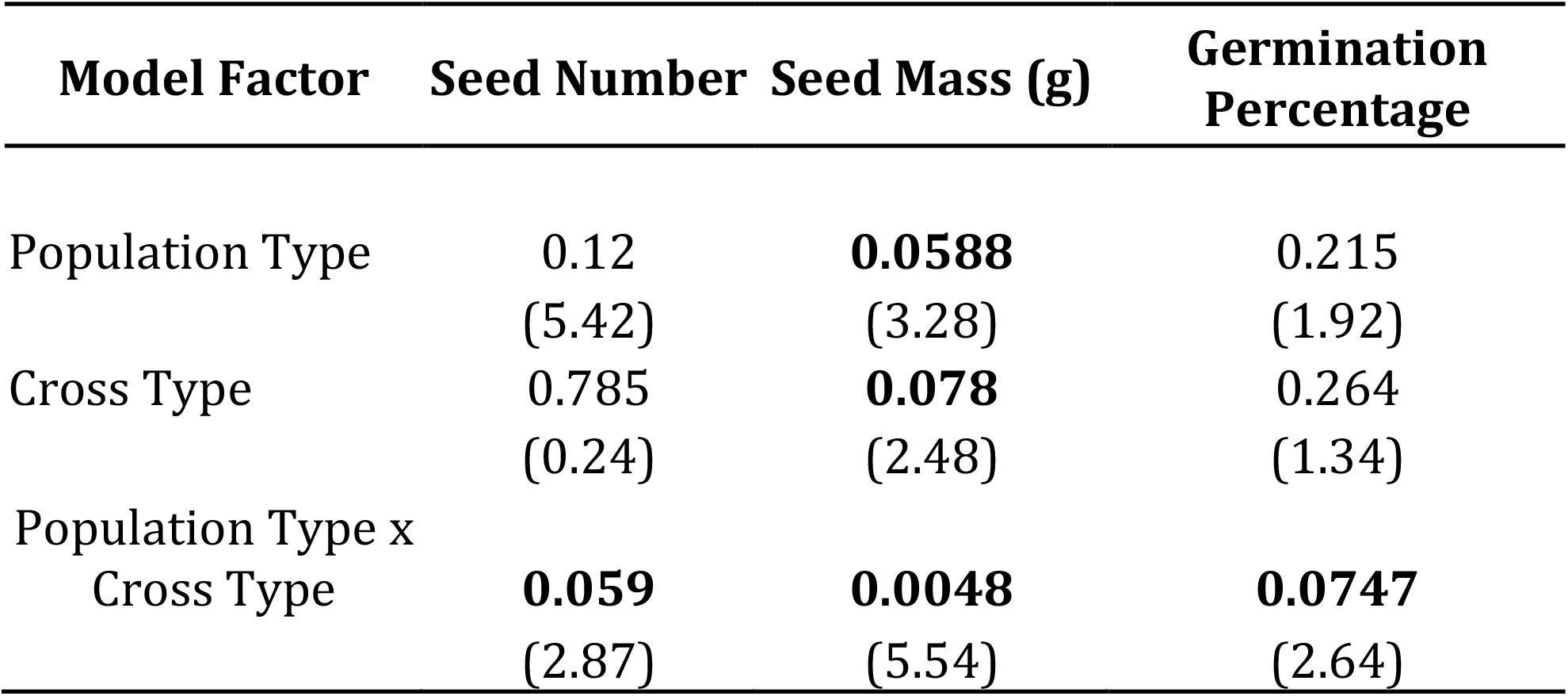
Results of a two-way ANOVA to determine the role of population type, crossing treatment, and a population type * crossing treatment interaction. P-values (*F*-value in parentheses) for each of the fixed effects are presented for each factor, seed number, seed mass, and germination percentage. Only seed mass showed a significant P-value < 0.5 for the interaction between population type and cross type, though a number of other factors showed nearly significant interaction effects, providing evidence for the role of population history in determining the evolutionary consequences of particular cross types.

| **Model Factor** | **Seed Number** | **Seed Mass (g)** | **Germination Percentage** |
| --- | --- | --- | --- |
| Population Type | 0.12<br>(5.42) | **0.0588**<br>(3.28) | 0.215<br>(1.92) |
| Cross Type | 0.785<br>(0.24) | **0.078**<br>(2.48) | 0.264<br>(1.34) |
| Population Type x<br>Cross Type | **0.059**<br>(2.87) | **0.0048**<br>(5.54) | **0.0747**<br>(2.64) |

**Table 4.** Means (and standard deviations) of seed, seedling, and vegetative traits of progeny from crosses in the greenhouse. Seed mass is the total weight of 50 seeds from a single capsule. Means for each population and P-values are provided in the following order: random within population cross, sib mating, and outcross. P-values represent the significance of the difference in mean between large and small populations for the separate cross types.

| Population Size | Seed Number | Seed Mass (g) | Germination Percentage |
| --- | --- | --- | --- |
| *a) Sib-matings* |  |  |  |
| Large, central populations | 218<br>(-96) | 0.0419<br>(-0.0101) | 74.43<br>(-16.78) |
| Small, isolated populations | 188<br>(-63) | 0.0404<br>(-0.0085) | 86.7<br>(-7.52) |
| **P-values** | **0.1355** | **0.3807** | **0.011** |
| *b) Within-population crosses* |  |  |  |
| Large, central populations | 218<br>(-105) | 0.0437<br>(-0.0109) | 73.39<br>(-17.4) |
| Small, isolated populations | 206<br>(-72) | 0.0377<br>(-0.0082) | 77.17<br>(-14.05) |
| **P-values** | **0.8246** | **0.0296** | **0.4603** |
| *c) Among-population crosses* |  |  |  |
| Large, central populations | 240<br>(-92) | 0.0444<br>(-0.01) | 79.7<br>(-15.3) |
| Small, isolated populations | 153<br>(-65) | 0.0323<br>(-0.0125) | 75.5<br>(-22.3) |
| **P-values** | **0.0011** | **0.0008** | **0.9029** |

There was significant inbreeding advantage for percent germination, and outbreeding depression for seed number, but only in small populations (Figure 3). More specifically, the progeny of sib-matings had a higher germination percentage than the progeny of within population crosses, and between population crosses produced a lower number of seeds than within population crosses (Figure 4). In large populations, there was no inbreeding depression and some evidence of heterosis, where in contrast to small populations, the progeny of between-population crosses had a higher germination percentage than the progeny of within-population crosses (Figure 4).

**Figure 3.**
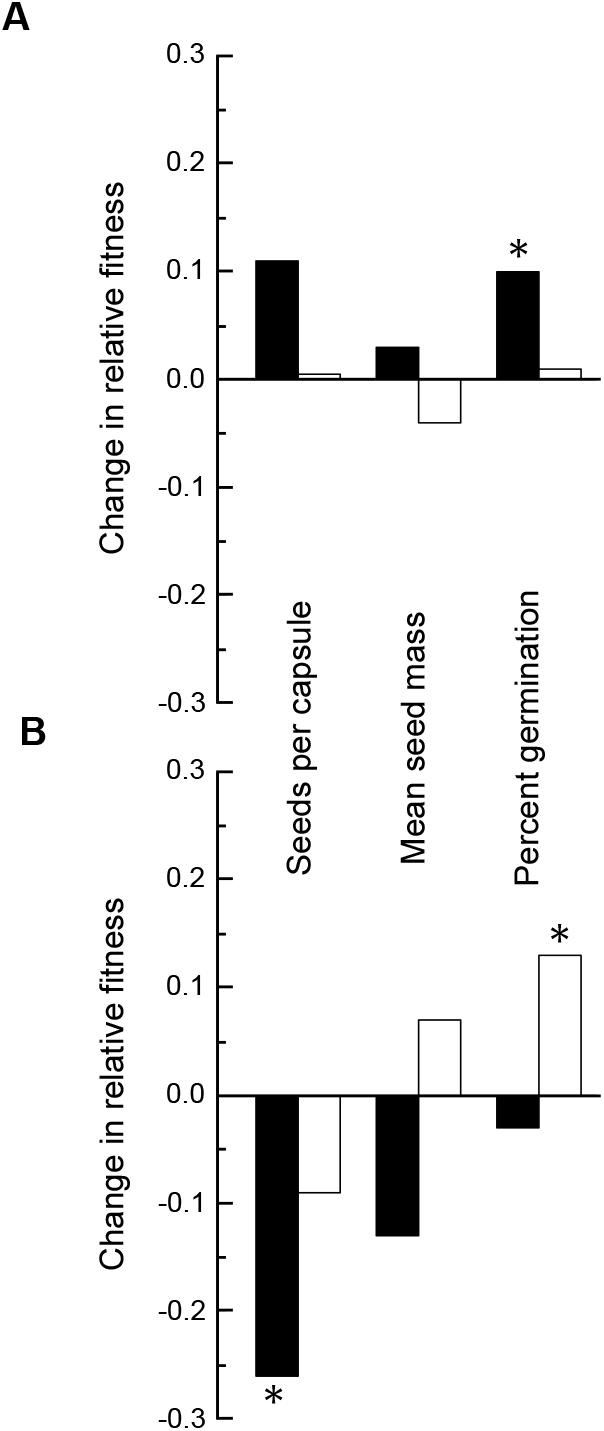
Differences in response to inbreeding and outcrossing for large (A) and small (B) populations. The horizontal baseline represents the average for each trait from the within population crosses. Open bars show the performance of progeny from crosses among full sibs relative to the within-population crosses; open bars above zero reflect an inbreeding advantage and open bars below zero reflect inbreeding depression. Solid bars show the performance of progeny from crosses between populations relative to the within-population crosses; Solid bars above zero indicate outcrossing advantage and solid bars below zero indicate outcrossing depression. Asterisks indicate that the two cross types used in that score were significantly different from each other (p<0.05).

**Figure 4.**
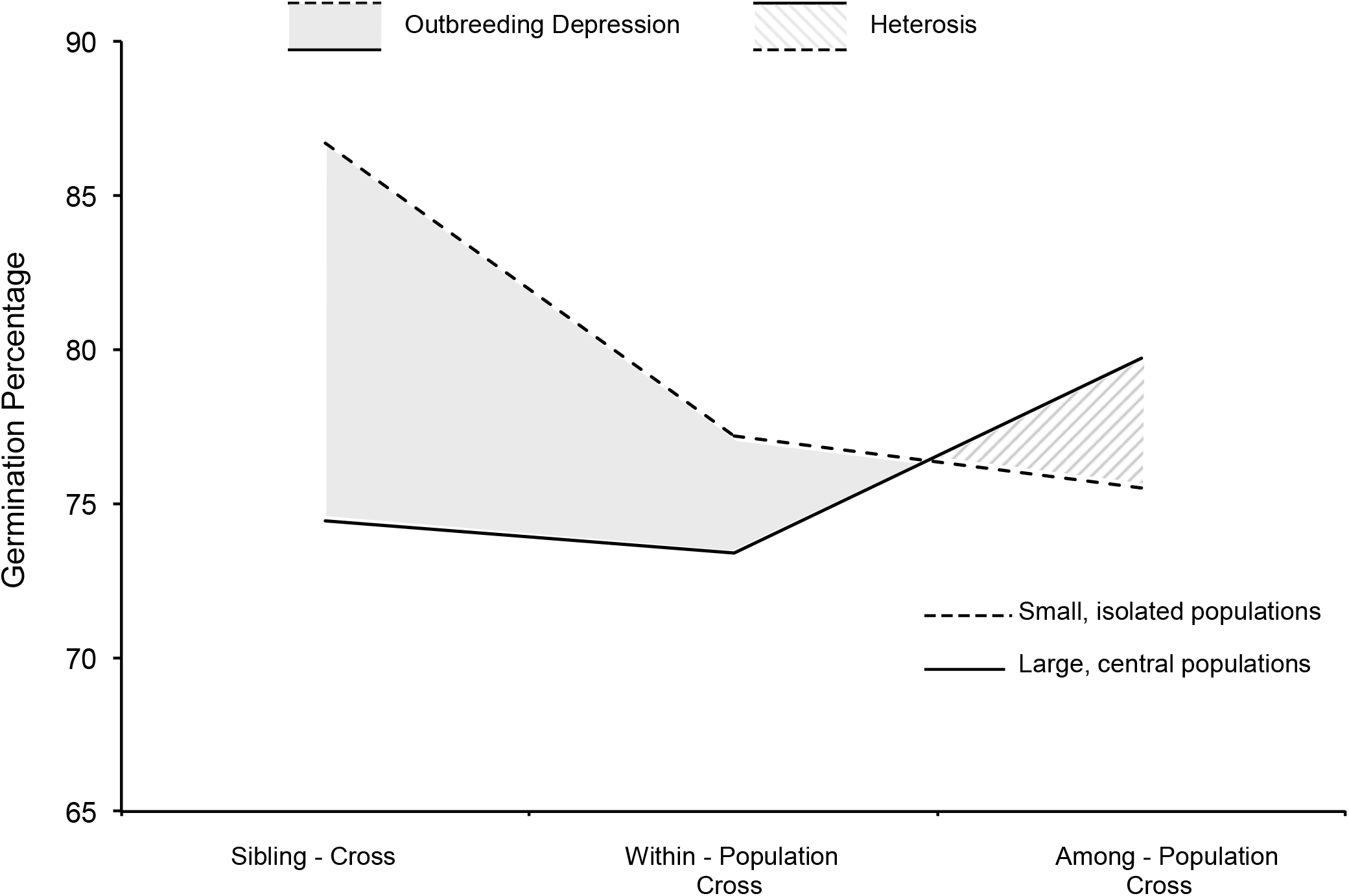
Percentage germination rate among sites within a cross type. Solid lines represent focal crosses from large, central populations; dashed lines represent crosses from small, isolated populations. In this representation, differences between types of crosses are indicated by the slope of the line, and asymmetries between types of populations results for each cross by different shadings.

## DISCUSSION

Our population genetic analysis suggests that populations vary significantly in the amount isolation they experience with spatial isolation and population size having substantive effects on the total amount of gene flow a population receives, as well as the subsequent relatedness of individuals available for breeding. In our study, the most surprising result was from the phenotypic characterization of plants and controlled crosses, where significant outbreeding depression was restricted to plants from small, isolated populations.

Outbreeding depression is believed to be common in angiosperms (Frankham 1995; Waser 1993) and is likely caused by the development of epistatic interactions within populations (Fenster et al. 1997). The finding that smaller, isolated populations exhibited more outbreeding depression therefore supports the prediction that founder effects will, in some cases, generate large shifts in allele frequencies (or allele combinations in the case of epistasis), potentially generating reproductive isolation and further genetic isolation between these and larger, central populations. In particular, the colonization process will aid in the generation of these complexes, consistent Wright’s shifting balances process (Wade 2013; Wright 1931; Wright 1932; Wright 1969; Wright 1977)

Population isolation has been hypothesized to increase inbreeding depression in the short-term, but given opportunities for subsequent purging, the long-term consequences of inbreeding on fitness become less predictable (Ives and Whitlock 2002; Whitlock 2002). Empirical evidence of purging under these, and perhaps most, conditions remains limited (Crnokrak and Barrett, 2002; though see Swindell and Bouzat, 2006; Facon, *et al*., 2011). Nonetheless, assuming theoretical predictions are correct, depending upon the magnitude of isolation and complex interactions with effective population size, we can expect an increase in variance in the consequences of inbreeding, and mean population fitness in general.

Metapopulation dynamics have been theorized to have substantive effects on the structuring of allelic variation (Wade and McCauley 1988; Whitlock and McCauley 1990). Population structure, as measured by *F*_ST_, has been theorized to have a significant effect on total genetic load by exposing recessive alleles to selection, possibly through dynamics such as (bi-parental-) inbreeding, an effect that is particularly apparent in the presence of hard selection (Whitlock 2002). Therefore, we can expect factors affecting *F*_ST_ to also play a large role in determining how particular controlled crosses, for example within-vs among-population crosses, affect fitness consequences (Agrawal, 2010; Agrawal and Whitlock, 2012). Disregarding these effects will lead to incorrect conclusions concerning the quantity of genetic load that segregates within a species.

Other studies have found individuals in small, isolated populations to have lower values of some fitness-related traits than individuals in large, central populations (Heschel and Paige 1995; Menges 1991), consistent with the results for seed count and seed mass in the present study. Seed mass has been shown to positively affect fitness in other plant species (Howell 1981; Kalisz 1986; Mazer and Schick 1991; Stanton 1985; Stanton 1984; Wulff 1986).

In our study, the results from small populations do not conform precisely to any of the predictions generated by simple models of inbreeding depression and the purging of deleterious recessive alleles, specifically a simple, inverse *F*-fitness relationship. Small, isolated populations had lower values for some traits related to fitness, as would be expected for populations experiencing inbreeding depression. However, three results from the crossing experiments contradict this generality: 1) progeny from among population crosses had lower seed production than progeny from within population crosses, 2) sib-mated progeny had a higher germination percentage than within population crosses, and 3) the only evidence of increased fitness of outcrossed progeny was in large populations, where outcrossing advantage was expected to be less pronounced. These results are reminiscent of a recent study using the perennial rosemary scrub (*Hypericum cumulicola*) where population size, age, and isolation had a significant effect on determining the fitness consequences of certain crosses (Oakley and Winn 2012). Oakley and Winn (2012) combined molecular marker-based estimators of migration and estimates of relative effective size for 16 natural populations of *H. cumulicola*. Fitness assays revealed that outbreeding advantage was significantly greater for effectively small populations relative to large ones, though there was an indication of outbreeding depression when comparing self-crosses to random-within population crosses.

The finding that outbreeding depression was significant only in small, isolated populations has several potential explanations. First, outbreeding depression can be the result of local adaptation (Fenster and Galloway 2000) and it is possible that local adaptation occurred more rapidly in isolated populations that are less likely to be inundated by gene flow from plants adapted to different environments (Antonovics 1968; Ellstrand and Elam 1993), consistent with the inverse relationship between isolation and resident proportion observed in the present study (Fig. 2). While this explanation is at least plausible for field-based observations of fitness, our controlled crosses took place in a controlled, greenhouse environment, and so this explanations provides little explanatory power here. Second some beneficial alleles may be partially recessive, and exposed to positive selection in more isolated population because those populations experience more (bi-parental-) inbreeding, wherein extreme population structure can be related to theoretical models of selfing species (Charlesworth, 1992; Hartfield and Glémin, 2014). However, given the persistently small population size, selection may remain ineffective on alleles providing both positive and negative fitness increments. Third, if there is significant epistatic variation segregating in larger populations, then genetic bottlenecks may fix certain combinations in local demes, leading to deleterious fitness consequences when those associations are disrupted during outcrossing (Corbett-Detig et al. 2013). Given we observed the presence of outbreeding depression within some populations this process may not always be effective, or at least not instantaneous, within demes. Finally, if colonizing propagules vary in their genetic load, selection may simply act in the long term on the persistence of individual lineages in their ability to persist in the absence of genetic rescue. Importantly, limited gene flow may then lead to, possibly ephemeral, outbreeding depression within these isolated populations.

The present results confirm that plants from large, central populations relative to small, isolated populations have differences in many traits related to fitness and that these differences have a genetic basis. However, as in previous studies, these results do not support a simple relationship between fitness and inbreeding, but instead support a growing body of evidence that the genetic and demographic outcomes of population structure likely generate variation in the *F*-fitness relationship. Thus, the metapopulation can be viewed as a heterogeneous landscape, with local variation in evolutionary outcomes. Importantly, these different evolutionary trajectories may influence the metapopulation ecology and vice versa, most obviously by influencing gene flow and population survival (Ingvarsson 2001; Newman and Pilson 1997; Saccheri et al. 1998). The connection between local evolutionary processes, and their effects on population growth and persistence, could also present a unique opportunity for studying the importance of inter-family or inter-demic selection (Schoen and Busch, 2008). From an applied perspective, our findings highlight the fact that inbreeding depression from deleterious recessive alleles is not the sole genetic mechanism affecting the long-term persistence of natural populations; genetic bottlenecks may have very different consequences across populations and through time.

## ACKNOWLEDGEMENTS

This work was supported by funding by the National Science Foundation (NSF) DEB #0919335 to Douglas Taylor and Janis Antonovics, and NSF-OISE# 1139716 to Douglas Taylor and Peter Fields. Joel Kniskern was supported by the Mountain Lake Biological Station NSF Research Experience for Undergraduates program.

## Notes

### Competing Interest Statement

The authors have declared no competing interest.

## LITERTURE CITED

Abdoullaye D, Acevedo I, Adebayo AA, Behrmann-Godel J, Benjamin RC, Bock DG, Born C, Brouat C, Caccone A, Cao LZ, et al. 2010 Permanent genetic rsources added to Molecular Ecology resources database 1 August 2009-30 September 2009. Mol Ecol Res. 10(1):232–236.

Agrawal AF. 2010 Ecological determinants of mutation load and inbreeding depression in subdivided populations. Am. Nat. 176(2):111–122.

Agrawal AF, Whitlock MC. 2012 Mutation load: the fitness of individuals in populations where deleterious alleles are abundant. In: Annual Review of Ecology, Evolution, and Systematics, Vol. 43: Annual Review of Ecology Evolution and Systematics (Futuyma DJ, ed), pp. 115–135. Annual Reviews, Palo Alto.

Antonovics J. 2004 Long term study of a plant-pathogen metapopulation. In: Ecology, genetics and evolution of metapopulations. (Hanski I, Gaggiotti O, eds), pp. 471–488. Academic Press.

Antonovics J, Thrall P, Jarosz A. 1998 Genetics and the spatial ecology of species interactions: the Silene-Ustilago system. In: Spatial ecology: the role of space in population dynamics and interspecific interactions. (Tilman D, Kareiva P, eds), pp. 158–180. Princeton University Press.

Antonovics J, Thrall P, Jarosz A, Stratton D. 1994 Ecological genetics of metapopulations: the Silene-Ustilago plant-pathogen system. In: Ecological genetics. (Real L, ed), pp. 146–170. Princeton University Press, NJ.

Baskin CC, Baskin JM. 1998 Seeds: ecology, biogeography, and evolution of dormancy and germination. Academic Press, San Diego, CA.

Byers DL, Waller DM. 1999 Do plant populations purge their genetic load? Effects of population size and mating history on inbreeding depression. Annu. Rev. Ecol. Syst. 30:479–513.

Caballero A. 1994 Developments in the prediction of effective population size. Heredity. 73(6):657–679.

Charlesworth B. 1992 Evolutionary rates in partially self-fertilizing species. Am. Nat. 140(1):126–148.

Charlesworth B. 2009 Effective population size and patterns of molecular evolution and variation. Nat Rev Genet. 10(3):195–205.

Charlesworth D. 2006 Evolution of plant breeding systems. Curr. Biol. 16(17):R726–R735.

Charlesworth D, Willis JH. 2009 The genetics of inbreeding depression. Nat Rev Genet. 10(11):783–796.

Corbett-Detig RB, Zhou J, Clark AG, Hartl DL, Ayroles JF. 2013 Genetic incompatibilities are widespread within species. Nature. 504(7478):135–137.

Crnokrak P, Barrett SCH. 2002 Perspective: Purging the genetic load: A review of the experimental evidence. Evolution. 56(12):2347–2358.

Dobzhansky T. 1937, 1941, 1951 Genetics and the Origin of Species. Columbia University Press, New York.

Dudash M, Carr D, Fenster C. 1997 Five generations of enforced selfing and outcrossing in Mimulus guttatus: Inbreeding depression variation at the population and family level. Evolution. 51(1):54–65.

Eppley SM, Pannell JR. 2009 Inbreeding depression in dioecious populations of the plant Mercurialis annua: comparisons between outcrossed progeny and the progeny of self-fertilized feminized males. Heredity. 102(6):600–608.

Escobar JS, Nicot A, David P. 2008 The different sources of variation in inbreeding depression, heterosis and outbreeding depression in a metapopulation of Physa acuta. Genetics. 180(3):1593–1608.

Facon B, Hufbauer Ruth a, Tayeh A, Loiseau A, Lombaert E, Vitalis R, Guillemaud T, Lundgren Jonathan, gEstoup A. 2011 Inbreeding Depression Is Purged in the Invasive Insect Harmonia axyridis. Curr. Biol. 21(5):424–427.

Gaggiotti OE, Bekkevold D, Jørgensen HBH, Foll M, Carvalho GR, Andre C, Ruzzante DE. 2009 Disentangling the effects of evolutionary, demographic, and environmental factors influencing genetic structure of natural populations: Atlantic Herring as a case study. Evolution. 63(11):2939–2951.

Goodnight, C. 1998 Epistasis and the effect of founder events on the additive genetic variance. Evolution 42(3):441–454

Grossenbacher D, Briscoe Runquist R, Goldberg EE, Brandvain Y. 2015 Geographic range size is predicted by plant mating system. Ecol. Lett. doi: 10.1111/ele.12449.n/a-n/a.

Guillaume F, Whitlock MC. 2007 Effects of migration on the genetic covariance matrix. Evolution. 61(10):2398–2409.

Hartfield M, Glémin S. 2014 Hitchhiking of deleterious alleles and the cost of adaptation in partially selfing species. Genetics. 196(1):281–293.

Heilbuth JC. 2000 Lower species richness in dioecious clades. Am. Nat. 156(3):221–241.

Ingvarsson P. 1998 Kin-structured colonization in Phalacrus substriatus. Heredity. 80:456–463.

Juillet N, Freymond H, Degen L, Goudet J. 2003 Isolation and characterization of highly polymorphic microsatellite loci in the bladder campion, Silene vulgaris (Caryophyllaceae). Mol Ecol Res. 3(3):358–359.

Keller LF, Waller DM. 2002 Inbreeding effects in wild populations. Trends Ecol. Evol. 17(5):230–241.

Keller SR, Gilbert KJ, Fields PD, Taylor DR. 2012 Bayesian inference of a complex invasion history revealed by nuclear and chloroplast genetic diversity in the colonizing plant, Silene latifolia. Mol Ecol. 21(19):4721–4734.

Matschiner M, Salzburger W. 2009 TANDEM: integrating automated allele binning into genetics and genomics workflows. Bioinformatics. 25(15):1982–1983.

Matute DR. 2013 The role of founder effects on the evolution of reproductive isolation. J. Evol. Biol. 26(11):2299–2311.

Mccauley DE. 1994 Intrademic group selection imposed by a parasitoid-host interaction. Am. Nat.:1–13.

Mccauley DE, Stevens J, Peroni P, Raveill J. 1996 The spatial distribution of chloroplast DNA and allozyme polymorphisms within a population of Silene alba (Caryophyllaceae). Am. J. Bot. 83(6):727–731.

Meirmans PG, Van Tienderen PH. 2004 GENOTYPE and GENODIVE: two programs for the analysis of genetic diversity of asexual organisms. Mol Ecol Notes. 4(4):792–794.

Moccia MD, Oger-Desfeux C, Marais GA, Widmer A. 2009 A white campion (Silene latifolia) floral expressed sequence tag (EST) library: annotation, EST-SSR characterization, transferability, and utility for comparative mapping. BMC Genomics. 10:243.

Moilanen A, Nieminen M. 2002 Simple connectivity measures in spatial ecology. Ecology. 83(4):1131–1145.

Mora MS, Mapelli FJ, Gaggiotti OE, Kittlein MJ, Lessa EP. 2010 Dispersal and population structure at different spatial scales in the subterranean rodent Ctenomys australis. BMC Genet. 11:14.

Muller HJ. 1942 Isolating mechanisms, evolution, and temperature. Biology Symposium. 6:71–125.

Pannell JR, Fields PD. 2014 Evolution in subdivided plant populations: concepts, recent advances and future directions. New Phytol. 201(2):417–432.

R Development Core Team. 2012. R: A language and environment for statistical computing. Vienna, Austria: R Foundation for Statistical Computing.

Rambaut A, Drummond AJ. 2009 Tracer v1.5 [Internet]. Available from: URL http://beast.bio.ed.ac.uk/Tracer.

Richards C. 2000 Inbreeding depression and genetic rescue in a plant metapopulation. Am. Nat. 155(3):383–394.

Richards C, Church S, Mccauley D. 1999 The Influence of Population Size and Isolation on Gene Flow by Pollen in Silene alba. Evolution. 53(1):63–73.

Schoen DJ, Busch JW. 2008 On the evolution of self-fertilization in a metapopulation. Int. J. Plant Sci. 169(1):119–127.

Swindell WR, Bouzat JL. 2006 Reduced inbreeding depression due to historical inbreeding in Drosophila melanogaster: evidence for purging. J. Evol. Biol. 19(4):1257–1264.

Teixeira S, Bernasconi G. 2007 High prevalence of multiple paternity within fruits in natural populations of Silene latifolia, as revealed by microsatellite DNA analysis. Mol Ecol. 16(20):4370–4379.

Thrall P, Richards C, Mccauley DE, Antonovics J. 1998 Metapopulation collapse: the consequences of limited gene flow in spatially structured populations. In: Modeling spatiotemporal dynamics in ecology. (Bascompte J, Sole RV, eds), pp. 83–104. Springer-Verlag Berlin.

Wheat CW, Fescemyer HW, Kvist J, Tas E, Vera JC, Frilander MJ, Hanski I, Marden JH. 2011 Functional genomics of life history variation in a butterfly metapopulation. Mol Ecol. 20(9):1813–1828.

Whitlock MC. 2002 Selection, load and inbreeding depression in a large metapopulation. Genetics. 160(3):1191–1202.

Whitlock MC, Ingvarsson PK, Hatfield T. 2000 Local drift load and the heterosis of interconnected populations. Heredity. 84(4):452–457.

Willi Y, Fischer M. 2005 Genetic rescue in interconnected populations of small and large size of the self-incompatible Ranunculus reptans. Heredity. 95(6):437–443.

Willi Y, Van Buskirk J, Fischer M. 2005 A threefold genetic allee effect: population size affects cross-compatibility, inbreeding depression and drift load in the self-incompatible Ranunculus reptans. Genetics. 169(4):2255–2265.

Wilson GA, Rannala B. 2003 Bayesian inference of recent migration rates using multilocus genotypes. Genetics. 163(3):1177–1191.

Wright S. 1938 Size of population and breeding structure in relation to evolution. Sci. 87:430–431.

Zhou S, Zhou C, Pannell JR. 2010 Genetic load, inbreeding depression and heterosis in an age-structured metapopulation. J. Evol. Biol.:2324–2332.

